# ORPA: A Fast and Efficient Method for Constructing Genome-Wide Alignments of Organelle Genomes for Phylogenetic Analysis

**DOI:** 10.1101/2023.05.26.542393

**Authors:** Guiqi Bi, Xinxin Luan, Jianbin Yan

**Author notes:** Correspondence to (G.B.); (J.Y.). These authors contributed equally to this work (Guiqi Bi, Xinxin Luan).

## Abstract

Creating a multi-gene alignment matrix for phylogenetic analysis using organelle genomes involves aligning single-gene datasets manually, a process that can be time-consuming and prone to errors. The HomBlocks pipeline has been created to eliminate the inaccuracies arising from manual operations. The processing of a large number of sequences, however, remains a time-consuming task. To conquer this challenge, we have developed a speedy and efficient method called ORPA. ORPA quickly generates multiple sequence alignments for whole-genome comparisons by parsing the result files of NCBI BLAST, completing the task in just one minute. With increasing data volume, ORPA’s efficiency is even more pronounced, over 300 times faster than HomBlocks in aligning 60 high-plant chloroplast genomes. The tool’s phylogenetic tree outputs demonstrate equivalent results to HomBlocks, indicating its outstanding efficiency. Due to its speed and accuracy, ORPA can identify species-level evolutionary conflicts, providing valuable insights into evolutionary cognition.

## INTRODUCTION

Phylogenetic trees utilizing organelle genomes are becoming indispensable in comparative genomics and systematics. They play a crucial role in elucidating the evolutionary relationships among species, particularly when incomplete lineage sorting obscures these relationships. This approach provides a broad perspective and facilitates a more accurate assessment of the phylogenetic relationships among species. It is a valuable tool for studying the complex evolutionary history of eukaryotic life (Li et al., 2019; Li et al., 2021).

Creating a precise multiple sequence alignment (MSA) is critical to constructing an accurate phylogenetic tree. This typically involves aligning single-copy genes beforehand and then concatenating them to form a super matrix. However, this can be laborious and error-prone, often requiring manual adjustments. To tackle this challenge, we previously introduced HomBlocks software (Bi et al., 2018), which enhances the accuracy and efficiency of MSA construction for organelle genome sequences by eliminating the need for manual operations.

However, when dealing with a substantial number of sequences, HomBlocks may still struggle with processing speed, which remains unsatisfactory. To address this issue more effectively, we have developed an innovative approach called ORPA. This method ranks as the fastest software for constructing multi-sequence alignments (MSA) of organelle genomes and delivers exceptionally accurate results compared to HomBlocks. Our research findings offer compelling evidence that ORPA is a highly viable option for HomBlocks, ensuring superior speed and accuracy. Moreover, the ORPA can be employed in systematic investigations to promptly obtain precise evolutionary relationships among species, resulting in significant research discoveries such as species-level evolutionary conflicts.

## RESULTS

### Comparison of tree topologies constructed from ORPA and HomBlocks sequence alignment

We evaluated the topological structures of phylogenetic trees constructed using ORPA and HomBlocks by employing the benchmark data used in the HomBlocks publication (Bi et al., 2018). As a direct comparison of the MSA results was impossible, we used this approach to observe similarities and differences between the two methods. The construction of phylogenetic trees was carried out using two approaches, the maximum likelihood method (RaxML (Stamatakis, 2014)) and Bayesian method (MrBayes 3.2.5 (Ronquist et al., 2012)).

In the first test dataset, we evaluated the performance of two software programs using a test dataset consisting of chloroplast genomes from 52 higher plants (Supplementary Table 1), as obtained from Zhang et al. (Zhang et al., 2016). Our results showed that the alignment lengths generated by ORPA and HomBlocks were 90,925 bp and 62,101 bp, respectively, with 8,270 bp and 8,404 bp of parsimony-informative sites. The resulting phylogenetic trees constructed from these two approaches exhibited identical topological structures with high support for all nodes except five (Fig. 2). For our second test dataset, we employed 36 mitochondrial genomes from Xenarthrans (Gibb et al., 2015) (Supplementary Table 2) to construct phylogenetic trees. Our comparative analysis of the resulting tree structures indicates that at non-100% support nodes, the support values from ORPA were slightly lower than those by HomBlocks. However, both methods exhibited a high degree of consistency in the overall topology of the phylogenetic tree (Fig. 3). These findings demonstrate the effectiveness of both approaches in generating reliable phylogenetic trees. Notably, all Multiple Sequence Alignment (MSA) constructions were accomplished within a time frame of less than 5 minutes using ORPA.

Moreover, we evaluated a set of higher plant mitochondrial datasets consisting of 18 publicly accessible mitochondrial sequences (Supplementary Table 3). Comparative assessments of the resulting phylogenetic trees corroborated the dependability of the ORPA and HomBlocks software suites (Fig. 3a). Additionally, we showcased the distribution of multiple sequence alignment (MSA) derived from both tools aligning to a reference genome, and the findings highlighted a remarkable concordance between the two methods, except for a few dissimilarities at specific loci (Fig. 3b).

### Comparing the Runtime of ORPA and HomBlocks

To directly compare the runtime differences of ORPA and HomBlocks on the same dataset, we tested 60 higher plant chloroplast genomes (Supplementary Table 4). First, we constructed genome-wide alignments using ORPA based on these 60 chloroplast sequences and conducted Maximum likelihood tree reconstruction using IQ-TREE (Nguyen et al., 2014). Then, we sampled based on the topology of the tree and compared the runtime of ORPA and HomBlocks. The sampling range increased by 5 with each deepening of the evolutionary relationship (Fig. 5a). We ran ORPA and HomBlocks for each dataset and calculated their respective running times in minutes. Since ORPA typically runs quickly, we standardized the comparison to 1 minute. Additionally, we used the alignment results from each dataset to construct a fast ML tree using IQ-TREE and compared the resulting trees generated by the two tools. To display the differences in running time, we presented a bar chart (Fig. 5b). For the comparison of the resulting trees from each dataset, we expressed the results as a similarity percentage (Fig. 5b).

Figure 5b shows a significant increase in HomBlocks’ runtime beyond 30 sequences, taking 313 minutes to complete the alignment process with 60 sequences. In contrast, based on the same data source, ORPA can process faster than HomBlocks when the number of sequences exceeds 10. Additionally, the runtime of HomBlocks increases exponentially with the number of sequences, which could become more significant with datasets containing over 60 sequences. Conversely, the script runtime for ORPA remains unaffected by the number of sequences, except during the data preparation stage. This is mainly attributed to the utilization of BLAST as the kernel for the alignment process, which avoids the need for single-threaded sequence comparison.

This highlights ORPA’s advantage over HomBlocks when dealing with a large number of sequences. Additionally, we conducted tree reconstructions for each pairwise test dataset and compared the ML tree topologies generated by ORPA and HomBlocks using treedist (https://github.com/agormp/treedist). Except for the comparison group with a sampling range of 25, the similarity between the tree topologies is 91%, indicating almost identical phylogenetic tree topologies generated by ORPA and HomBlocks in the other 11 comparison groups. Our findings suggest that ORPA outperforms HomBlocks in terms of speed and accuracy, offering researchers a powerful tool for creating whole-genome alignments.

### Using ORPA for rapid detection of systematic evolutionary conflicts

Advancements in sequencing technology have led to the accumulation of a vast amount of organellar genome data. Effective utilization of this data has become a growing field of interest. This is especially important for newly sequenced data, as rapid confirmation of species’ evolutionary relationships is crucial to verify sequencing accuracy. Additionally, constructing phylogenetic trees with speed and accuracy to investigate evolutionary conflicts is a key area of research in systematics biology. ORPA offers an elegant approach to achieving these goals. To provide a more comprehensive demonstration, we utilized 52 Lamiales chloroplast datasets as an example (Supplementary Table 5). We employed ORPA to build a multiple sequence alignment of 101,454 characters, and constructed a corresponding phylogenetic tree to depict the evolutionary relationships. Figure 5 demonstrates that there are two apparent conflicts between the phylogenetic branches in regards to *Wightia speciosissima* and *Comoranthus minor*.

*Wightia speciosissima*, an angiosperm, has been assigned to a distinct family (Wightiaceae) by the Angiosperm Phylogeny Group IV (The Angiosperm Phylogeny, 2016). Its previous classification placed it within the Paulowniaceae family. However, analysis of its evolutionary branching suggests that it shares closer evolutionary relationships with the Phrymaceae family, thus representing a distinct lineage. This observation was also made by Xia et al (Xia et al., 2019). Their study on plant phylogeny utilized data from nine chloroplast genes and one mitochondrial *rps*3 gene. Consequently, they advise against including *Wightia speciosissima* in the Paulowniaceae family and suggest that it may instead be a hybrid origin between early lineages of Phrymaceae and Paulowniaceae (Xia et al., 2019).

This phylogenetic tree also reveals the incongruity between the genus *Comoranthus* and *Schrebera* in terms of their phylogenetic relationships. *Schrebera* is found in Africa and India, while *Comoranthus* is only found in Madagascar and the Comoros Islands (Wallander et al., 2000). Both species have similar fruit morphology: capsules with a woody ovary, ruffled epidermis, and split in half when mature. They contain seeds and are suborbicular and pear-shaped (Engel, 1968). Our analysis of the evolutionary relationships among these studied species reveals a paraphyletic relationship, with *Comoranthus* found nested within *Schrebera*. This outcome is consistent with recent findings by Hong-Wa et al., who suggest the genera should be synonymized (Hong-Wa et al., 2023). Incorporating this finding into taxonomic classification will aid in a more accurate understanding of the evolutionary history of these plant groups.

In summary, ORPA has demonstrated considerable promise in the realm of systematic taxonomy, as demonstrated by the aforementioned use cases.

## EXPERIMENTAL PROCEDURES

### Methods

The framework of ORPA is written in Perl. As tools for aligning genomes, HomBlocks uses a method of identifying locally collinear blocks (LCBs), while the main difference with ORPA is its strategy of directly parsing the NCBI BLAST online tool results. By avoiding the need for software installation and various dependencies, this approach simplifies genome alignment for novices in the field of bioinformatics (Fig. 1). The core of ORPA is based on the widely-used BLAST tool (McGinnis et al., 2004), which offers significant improvements in the efficiency and speed of sequence alignments. Compared to HomBlocks, ORPA is able to construct alignment files within 5 minutes on average. In contrast, HomBlocks requires an increasing amount of processing time as the number of sequences being aligned grows due to the single-threaded operation of its core software, Mavue (Darling et al., 2004). Therefore, ORPA offers a more efficient and versatile alternative to HomBlocks.

**Fig. 1.**
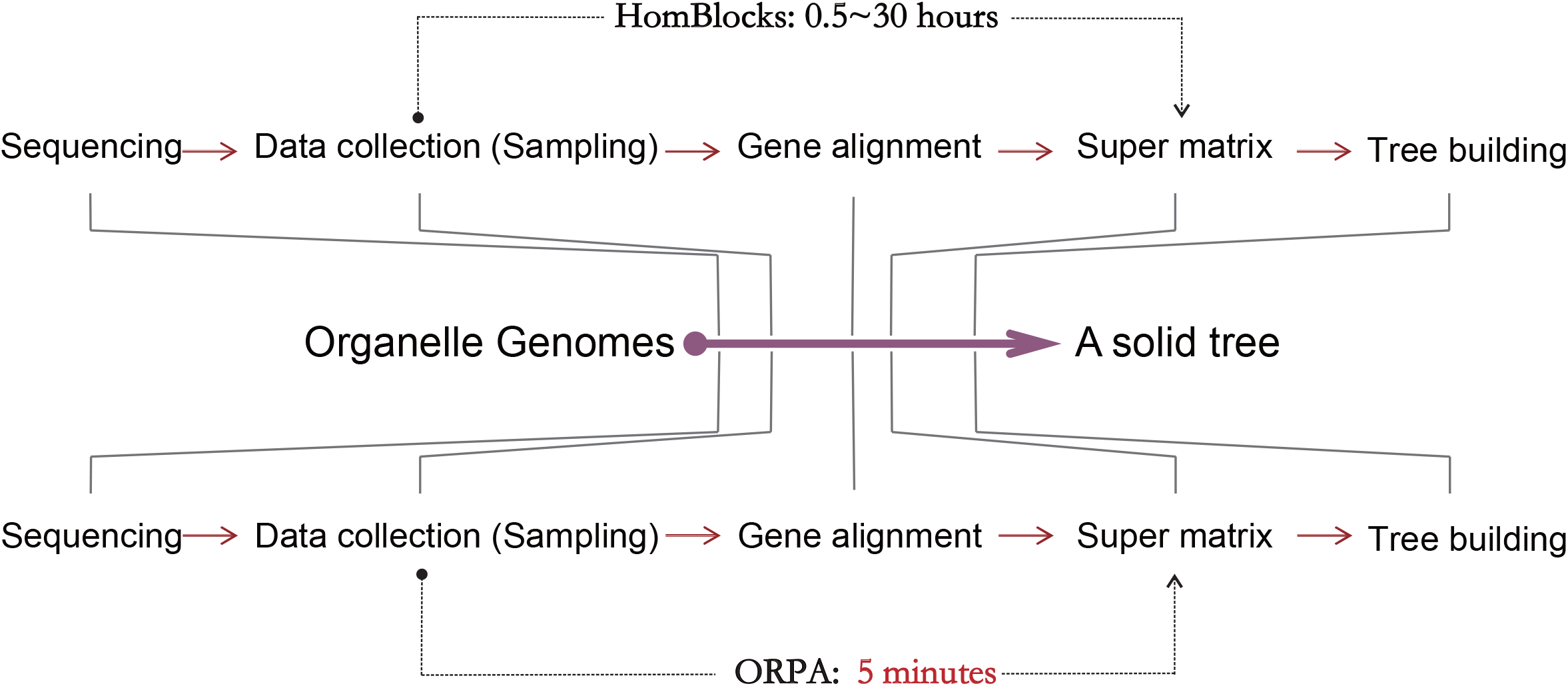
Comparison of ORPA and HomBlocks efficiency in the conventional workflow for the phylogenetic tree construction of organelle genomes.

**Fig. 2.**
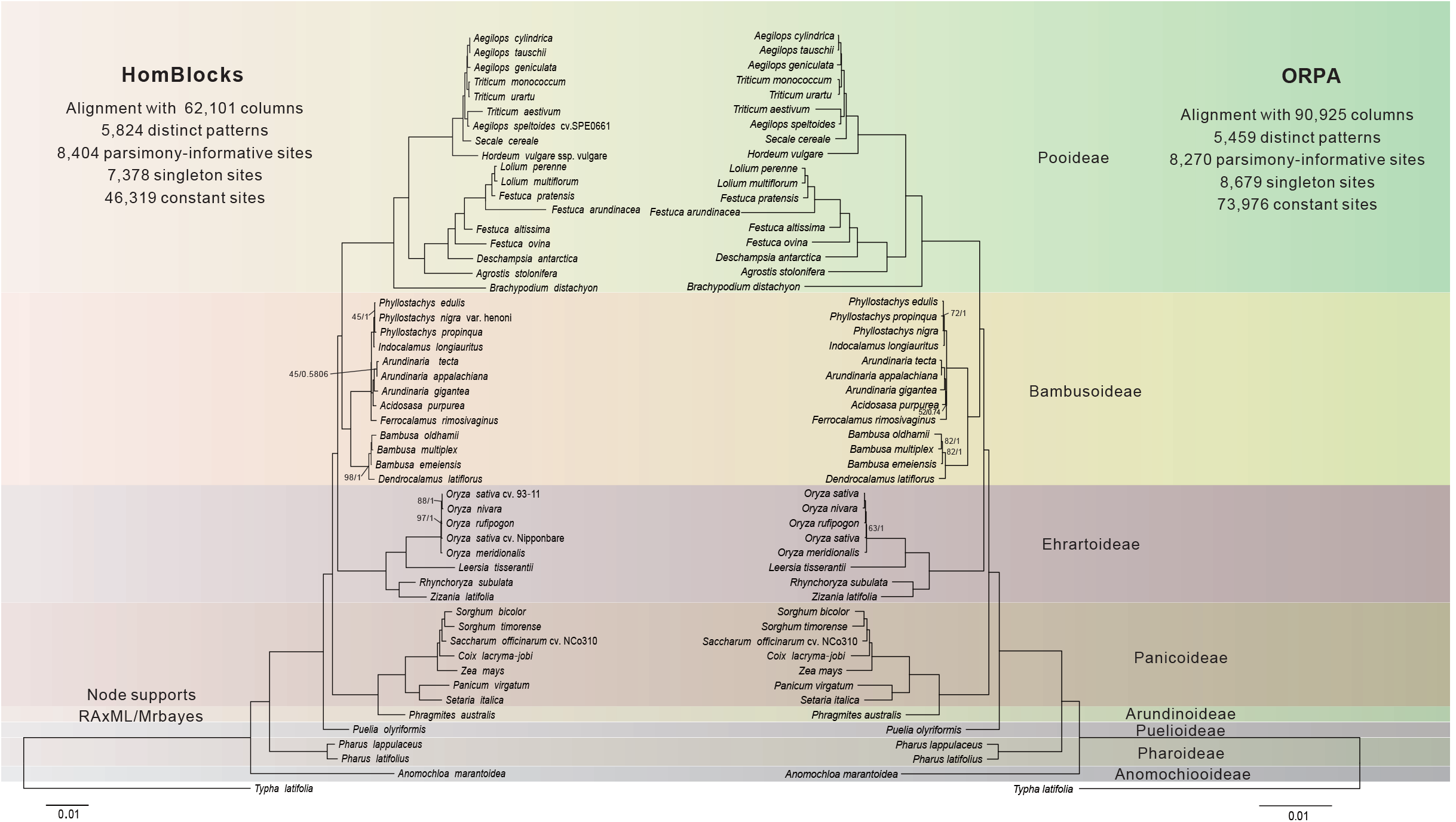
Comparison of topology between the HomBlocks tree (left) and the ORPA tree (right) of 52 higher plant chloroplast genomes. The phylogenetic trees were constructed using maximum likelihood (ML) and Bayesian inference (BI) methods with the HomBlocks alignment (62,101 characters) and the ORPA alignment (90,925 characters), respectively. The support values inferred from RAxML (left) and Bayesian posterior probability (right) are indicated by the numbers on the nodes. Fully resolved nodes are not labeled with numbers. These results provide insights into the comparative performance of the two alignment methods for phylogenetic analysis of chloroplast genomes in higher plants.

**Fig. 3.**
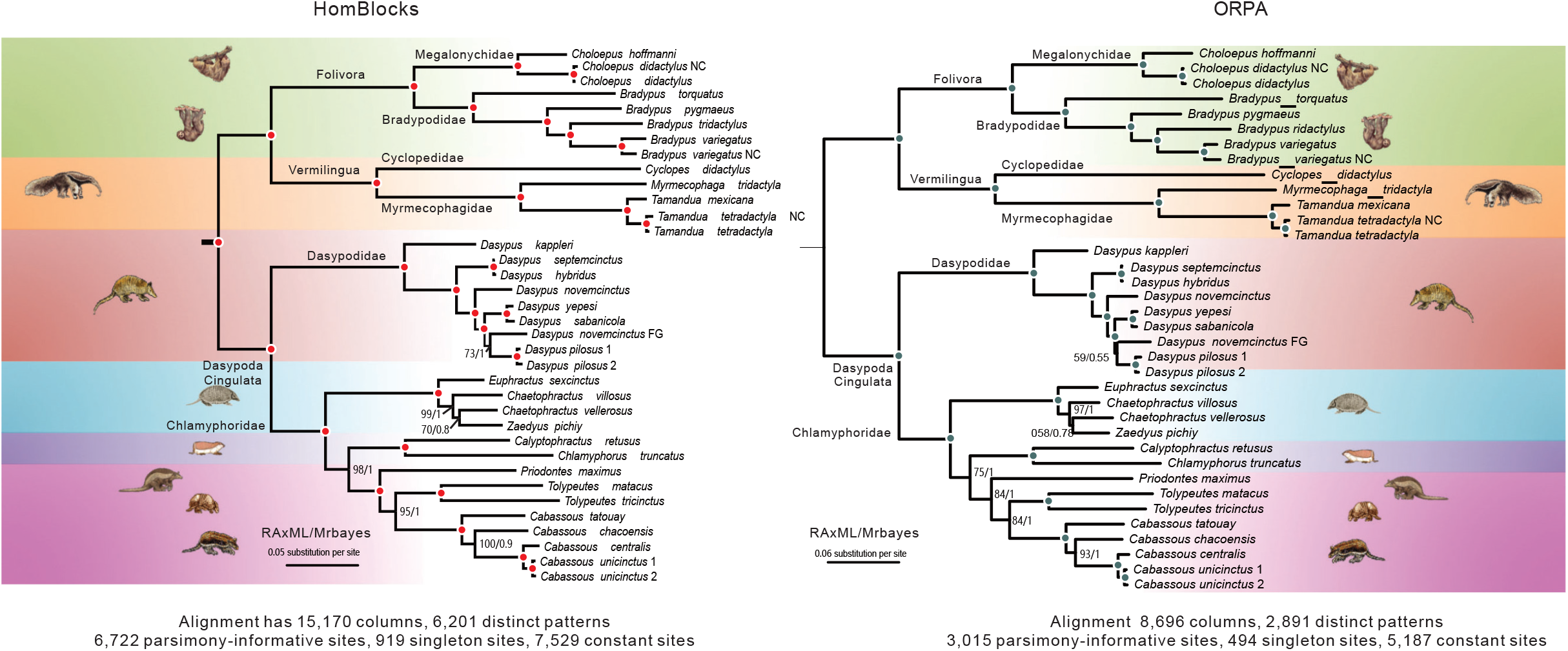
Topology comparison of two phylogenetic trees of 36 xenarthran mitochondrial genomes. The HomBlocks tree (left) and the ORPA tree (right) were constructed using different alignment methods, one with 15,170 characters and the other with 8,696 characters. Maximum likelihood and Bayesian inference methods were used to construct the trees, and the support values derived from RAxML (left) and Bayesian posterior probability (right) are indicated by the numerical values on the nodes. Fully resolved nodes are unlabeled.

**Fig. 4.**
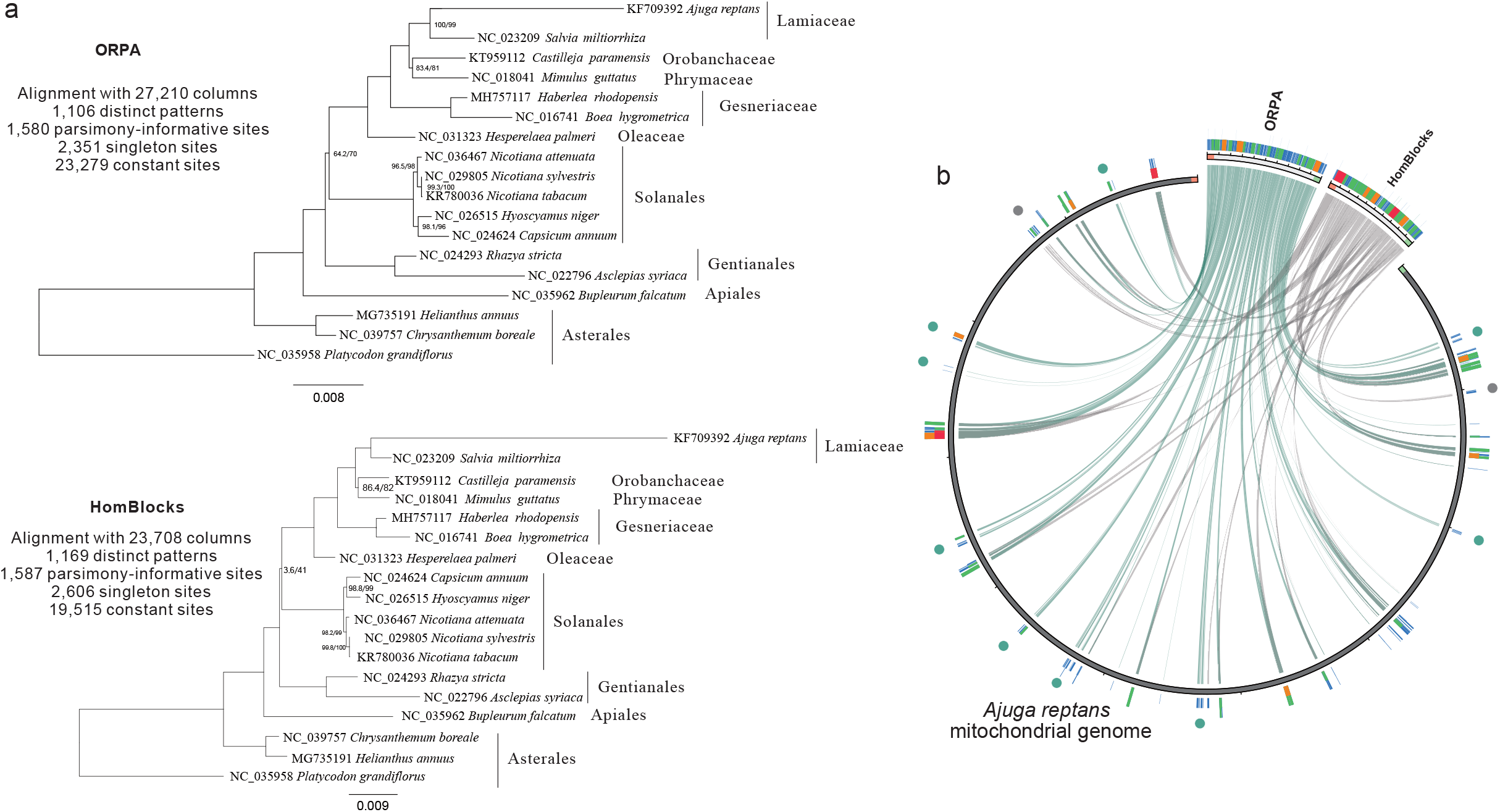
Comparison of phylogenetic trees and alignment methods for 18 higher plant mitochondrial genomes. **a**, Two phylogenetic trees of 18 higher plant mitochondrial genomes were constructed using ORPA and HomBlocks alignment methods, respectively. The trees were constructed with maximum likelihood and Bayesian inference methods, and the support values derived from RAxML and Bayesian posterior probability are indicated on each node. Fully resolved nodes are unlabeled. **b**, Distributional differences of phylogenetic alignments obtained from ORPA and HomBlocks methods using Ajuga reptans as the reference sequence. The circos plot illustrates the differing sequence composition sites between the two methods, with green and gray dots indicating the variation between the alignments.

**Fig. 5.**
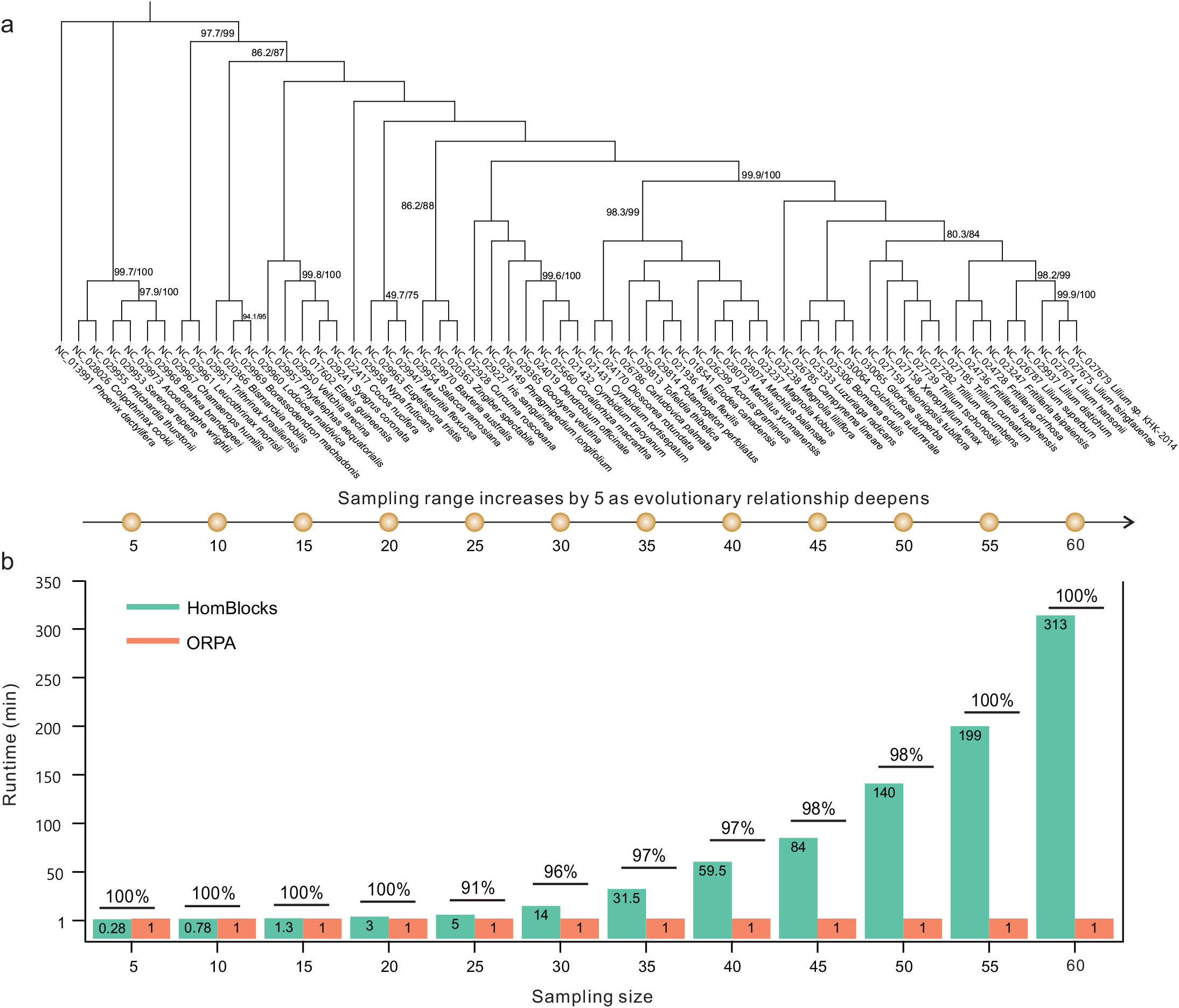
Comparison of ORPA and HomBlocks runtime efficiency. **a**, Comparison of runtime for 60 higher plant chloroplast datasets. A maximum likelihood tree shows the evolutionary relationship among 60 samples. Nodes with 100% support are unspecified, and other partially supported nodes are labeled with bootstrap and aLTR values. Sampling begins at the base of the tree and proceeds with increasing sample sizes of 5 until all data are used, resulting in a total of 12 comparison groups. **b**, Comparison of ORPA and HomBlocks runtime. The sample size corresponds to the sampling range in Figure 5a. The percentage on the bar chart represents the similarity in systemic tree topology generated by the two software programs.

**Fig. 6.**
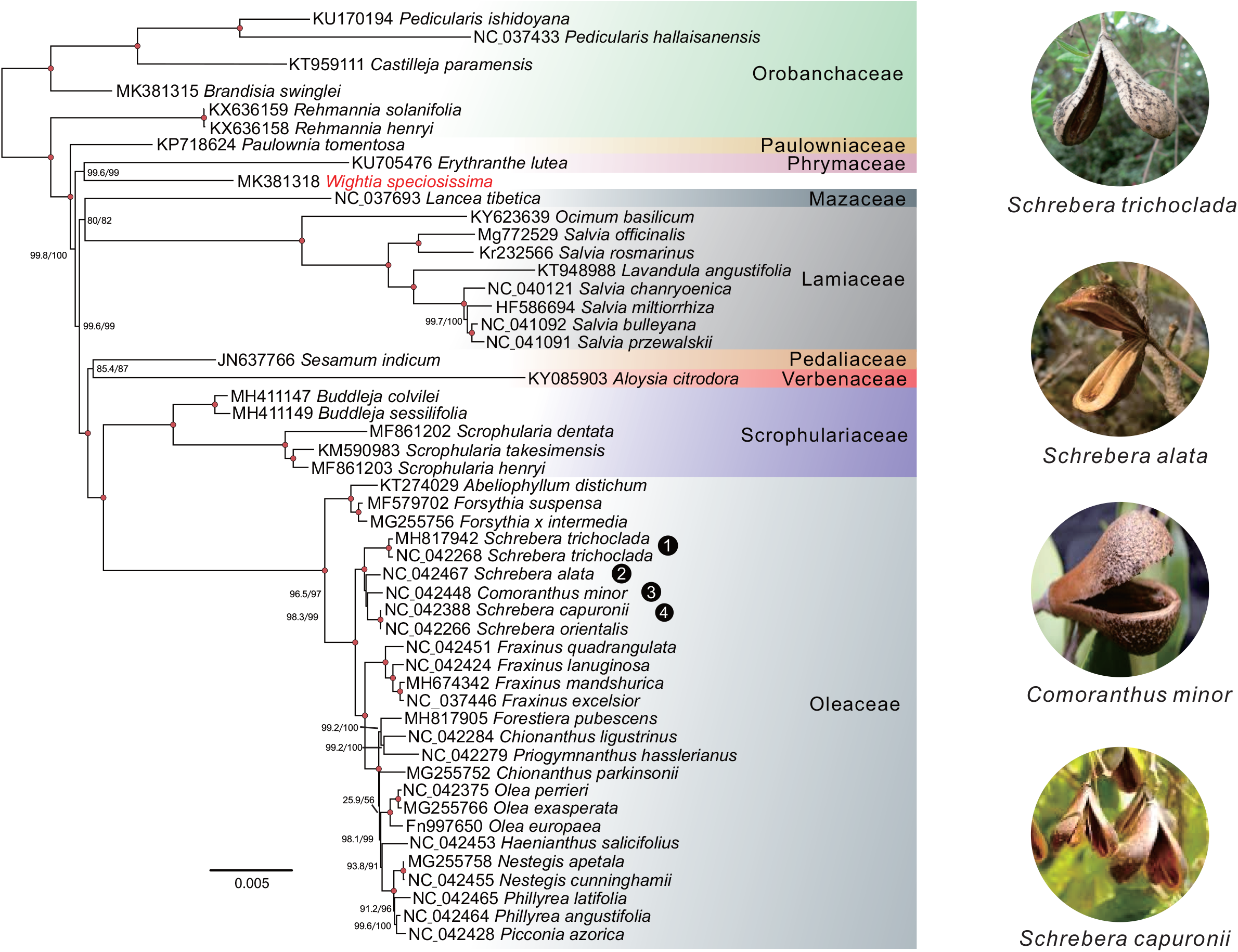
Identification of species-level evolutionary conflicts using ORPA. A total of 52 Lamiales chloroplast trees were constructed using 101,544 characters from the ORPA alignment. Maximum likelihood and Bayesian inference methods were used to construct the trees, and the support values derived from RAxML (left) and Bayesian posterior probability (right) are indicated by numerical values on the nodes. Fully resolved nodes are indicated by red dots. *Wightia speciosissima*, which has a controversial position in Lamiales, is labelled in red. The morphology of four species from *Schrebera* and *Comoranthus* genera is shown on the right side of the figure. Additionally, the results reveal a paraphyletic relationship, with *Comoranthus minor* nested within *Schrebera*, leading to the synonymization of these genera.

ORPA also provides users with four trimming methods, namely Gblocks (Castresana, 2000), trimAl (Capella□Gutiérrez et al., 2009), Noisy (Dress et al., 2008), and BMGE (Criscuolo et al., 2010), which are same to those offered by HomBlocks. Importantly, users can directly use the output results from ORPA to facilitate the construction of a phylogenetic tree, thus streamlining the sequence alignment process.

### Implementation

ORPA is a rapid tool for constructing multiple sequence alignments of organelles. It is a command-line tool and functional under any version of Linux without the need for external installation. The Perl source code of ORPA is freely available for download at https://github.com/BGQ/ORPA.git, and comprehensive documentation and tutorials can be found at https://github.com/BGQ/ORPA.git.

## ACKNOWLEDGMENTS

We sincerely thank the editors and reviewers for their valuable suggestions and comments on this study. This work was supported by the National Key R&D Program of China (2018YFA0903200), Science Technology and Innovation Commission of Shenzhen Municipality of China (ZDSYS 20200811142605017). It was also supported by the Elite Young Scientists Program of CAAS and the Agricultural Science and Technology Innovation Program.

## AUTHOR CONTRIBUTIONS

GQB and JBY conceived and designed the study. GQB and XXL collected data. GQB and XXL performed data analysis. GQB and JBY wrote the manuscript with other authors providing recommendations for modifications.

## DATA AVAILABILITY STATEMENT

The attached file contains a list of example data and organelle genome sequences that are referred to in the main body of this study.

## SUPPORTING INFORMATION

Additional Supporting Information may be found in the online version of this article.

**Table S1**. Accession numbers of 52 higher plant chloroplast genomes from GenBank used in the first example dataset.

**Table S2**. Accession numbers of 36 xenarthrans mitochondrial genomes from GenBank used in the second example dataset.

**Table S3**. Accession numbers of 18 higher plant mitochondrial genomes from GenBank used in the third example dataset.

**Table S4**. Accession numbers of 60 higher plant chloroplast genomes from GenBank used in the fourth example dataset.

**Table S5**. Accession numbers of Lamiales chloroplast genomes from GenBank used in the fifth example dataset.

