## Supplementary information for "ORPA: A Fast and Efficient Method for Constructing Genome-Wide Alignments of Organelle Genomes for Phylogenetic Analysis"

**Table S1. Species of 52 higher plant chloroplast genomes utilized in the first example dataset and their corresponding accession No. in GenBank database.**

| **Species** | **Length** | **GC content** | **Accession NO.** |
| --- | --- | --- | --- |
| *Acidosasa purpurea* | 139,697 bp | 38.90% | HQ337793.1 |
| *Aegilops cylindrica* | 113,490 bp | 37.29% | NC_023096.1 |
| *Aegilops geniculata* | 113,893 bp | 37.25% | NC_023097.1 |
| *Aegilops speltoides* isolate SPE0661 | 113,536 bp | 37.25% | JQ740834.1 |
| *Aegilops tauschii* | 114,112 bp | 37.26% | NC_022133.1 |
| *Agrostis stolonifera* | 136,584 bp | 38.45% | NC_008591.1 |
| *Anomochloa marantoidea* | 138,412 bp | 38.66% | GQ329703.1 |
| *Arundinaria appalachiana* | 139,547 bp | 38.91% | NC_023934.1 |
| *Arundinaria gigantea* | 138,935 bp | 38.93% | NC_020341.1 |
| *Arundinaria tecta* | 139,499 bp | 38.92% | NC_023935.1 |
| *Bambusa emeiensis* | 139,493 bp | 38.91% | HQ337797.1 |
| *Bambusa multiplex* | 139,394 bp | 38.91% | NC_024668.1 |
| *Bambusa oldhamii* | 139,350 bp | 38.92% | FJ970915.1 |
| *Brachypodium distachyon* | 135,199 bp | 38.57% | NC_011032.1 |
| *Coix lacryma-jobi* | 140,745 bp | 38.46% | FJ261955.1 |
| *Dendrocalamus latiflorus* | 139,394 bp | 38.92% | FJ970916.1 |
| *Deschampsia antarctica* | 135,362 bp | 38.32% | KF887484.1 |
| *Ferrocalamus rimosivaginus* | 139,467 bp | 38.86% | HQ337794.1 |
| *Festuca altissima* | 135,272 bp | 38.45% | JX871939.1 |
| *Festuca arundinacea* | 136,048 bp | 38.39% | NC_011713.2 |
| *Festuca ovina* | 133,165 bp | 38.37% | JX871940.1 |
| *Festuca pratensis* | 135,291 bp | 38.27% | JX871941.1 |
| *Hordeum vulgare subsp. vulgare* | 136,462 bp | 38.32% | NC_008590.1 |
| *Indocalamus longiauritus* | 139,668 bp | 38.91% | HQ337795.1 |
| *Leersia tisserantii* | 1365,51 bp | 38.88% | JN415112.1 |
| *Lolium multiflorum* | 135,175 bp | 38.25% | JX871942.1 |
| *Lolium perenne* | 135,282 bp | 38.25% | NC_009950.1 |
| *Oryza meridionalis* | 134,558 bp | 39.01% | NC_016927.1 |
| *Oryza nivara* | 134,494 bp | 39.01% | NC_005973.1 |
| *Oryza rufipogon* | 134,544 bp | 39.00% | NC_017835.1 |
| *Oryza sativa* Indica Group isolate 93-11 | 134,496 bp | 39.00% | AY522329.1 |
| *Oryza sativa* Japonica Group cultivar Nipponbare | 134,551 bp | 39.00% | KM088016.1 |
| *Panicum virgatum* | 139,619 bp | 38.60% | NC_015990.1 |
| *Pharus lappulaceus* | 141,928 bp | 38.43% | NC_023245.1 |
| *Pharus latifolius* | 142,077 bp | 38.37% | JN032131.1 |
| *Phragmites australis* | 137,614 bp | 38.66% | KJ825856.1 |
| *Phyllostachys edulis* | 139,679 bp | 38.88% | HQ337796.1 |
| *Phyllostachys nigra var. henonis* | 139,839 bp | 38.90% | HQ154129.1 |
| *Phyllostachys propinqua* | 139,704 bp | 38.88% | JN415113.1 |
| *Puelia olyriformis* | 140,344 bp | 38.97% | NC_023449.1 |
| *Rhynchoryza subulata* | 136,303 bp | 39.00% | JN415114.1 |
| *Saccharum hybrid* cultivar NCo 310 | 141,182 bp | 38.44% | NC_006084.1 |
| *Secale cereale* | 114,843 bp | 37.20% | NC_021761.1 |
| *Setaria italica* | 138,833 bp | 38.62% | KJ001642.1 |
| *Sorghum bicolor* | 140,754 bp | 38.49% | NC_008602.1 |
| *Sorghum timorense* | 140,629 bp | 38.50% | KF998272.1 |
| *Triticum aestivum* | 134,545 bp | 38.31% | NC_002762.1 |
| *Triticum monococcum* | 116,399 bp | 37.37% | NC_021760.1 |
| *Triticum urartu* | 115,773 bp | 37.38% | NC_021762.1 |
| *Typha latifolia* | 161,572 bp | 36.61% | GU195652.1 |
| *Zea mays* | 140,384 bp | 38.46% | NC_001666.2 |
| *Zizania latifolia* | 136,461 bp | 39.00% | NC_029401.1 |

**Table S2. Species of 36 *xenarthrans* mitochondrial genomes utilized in the second example dataset and their corresponding accession No. in GenBank database.**

| **Species** | **Length** | **GC content** | **Accession NO.** |
| --- | --- | --- | --- |
| *Bradypus pygmaeus* voucher USNM_579179 | 16,799 bp | 44.10% | NC_028554.1 |
| *Bradypus torquatus* voucher BA449/11 | 16,714 bp | 41.88% | NC_028555.1 |
| *Bradypus tridactylus* voucher ISEM_T-5013 | 16,920 bp | 43.98% | KT818525.1 |
| *Bradypus variegatus* voucher MVZ_155186 | 16,641 bp | 44.35% | KT818526.1 |
| *Bradypus variegatus* | 16,975 bp | 44.26% | NC_006923.1 |
| *Cabassous centralis* voucher AMNH_MO-10752 | 17,114 bp | 42.80% | NC_028556.1 |
| *Cabassous chacoensis* voucher ISEM_T-2350 | 16,280 bp | 42.89% | NC_028557.1 |
| *Cabassous tatouay* voucher ZVC_M365 | 16,638 bp | 43.33% | NC_028558.1 |
| *Cabassous unicinctus* voucher ISEM_T-2291 | 16,397 bp | 43.12% | NC_028559.1 |
| *Cabassous unicinctus* voucher MNHN_1999-1068 | 16,468 bp | 43.14% | KT818531.1 |
| *Calyptophractus retusus* voucher ZSM_T-Bret | 16,231 bp | 44.70% | KT818532.1 |
| *Chaetophractus vellerosus* voucher ISEM_T-CV1 | 16,598 bp | 40.23% | NC_028561.1 |
| *Chaetophractus villosus* voucher ISEM_T-NP390 | 16,796 bp | 39.77% | NC_028562.1 |
| *Chlamyphorus truncatus* voucher ISEM_T-CT1 | 16,245 bp | 44.15% | NC_028563.1 |
| *Choloepus didactylus* voucher MNHN_1998-1819 | 16,487 bp | 43.30% | KT818537.1 |
| *Choloepus didactylus* | 16,543 bp | 43.26% | AY960980.1 |
| *Choloepus hoffmanni* voucher ISEM_T-6052 | 16,386 bp | 42.96% | KT818538.1 |
| *Cyclopes didactylus* voucher MNHN_1998-234 | 16,368 bp | 36.88% | NC_028564.1 |
| *Dasypus hybridus* voucher ZVC_M2010 | 17,013 bp | 38.92% | NC_028565.1 |
| *Dasypus kappleri voucher ISEM_T-3365* | 16,636 bp | 41.34% | KT818541.1 |
| *Dasypus novemcinctus* voucher ISEM_T-1863 | 16,487 bp | 39.61% | KT818542.1 |
| *Dasypus novemcinctus* | 17,056 bp | 38.86% | Y11832.1 |
| *Dasypus pilosus* voucher LSUMZ_21888 | 17,050 bp | 38.97% | KT818543.1 |
| *Dasypus pilosus* voucher MSB_49990 | 16,999 bp | 38.83% | KT818544.1 |
| *Dasypus sabanicola* voucher USNM_372834 | 16,936 bp | 38.93% | NC_028568.1 |
| *Dasypus septemcinctus* voucher ISEM_T-3002 | 16,683 bp | 39.32% | NC_028569.1 |
| *Dasypus yepesi voucher* MLP_30.III.90.2 | 16,917 bp | 38.93% | NC_028570.1 |
| *Euphractus sexcinctus* voucher ISEM_T-1246 | 16,602 bp | 40.29% | NC_028571.1 |
| *Myrmecophaga tridactyla* voucher ISEM_T-2862 | 16,498 bp | 39.10% | NC_028572.1 |
| *Priodontes maximus* voucher ISEM_T-2353 | 16,492 bp | 44.44% | NC_028573.1 |
| *Tamandua mexicana* voucher MVZ_192699 | 16,385 bp | 38.94% | NC_028574.1 |
| *Tamandua tetradactyla* voucher ISEM_T-6054 | 16,393 bp | 39.07% | KT818552.1 |
| *Tamandua tetradactyla* | 16,395 bp | 39.08% | AJ421450.1 |
| *Tolypeutes matacus* voucher ISEM_T-2348 | 16,980 bp | 40.97% | NC_028575.1 |
| *Tolypeutes tricinctus* voucher JB21 | 17,161 bp | 41.35% | NC_028576.1 |
| *Zaedyus pichiy voucher* ISEM_T-6060 | 16,598 bp | 40.17% | NC_028577.1 |

**Table S3. Species of 18 higher plant mitochondrial genome utilized in the third example dataset and their corresponding accession No. in GenBank database.**

| **Species** | **Length** | **GC content** | **Accession NO.** |
| --- | --- | --- | --- |
| *Ajuga reptans* | 352,069bp | 45.10% | KF709392 |
| *Nicotiana tabacum* | 430,863bp | 44.95% | KR780036 |
| *Castilleja paramensis* | 495,499bp | 43.52% | KT959112 |
| *Helianthus annuus* | 305,217bp | 45.05% | MG735191 |
| *Haberlea rhodopensis* | 484,138bp | 44.10% | MH757117 |
| *Boea hygrometrica* | 510,519bp | 43.27% | NC_016741 |
| *Mimulus guttatus* | 525,671bp | 45.14% | NC_018041 |
| *Asclepias syriaca* | 682,498bp | 43.43% | NC_022796 |
| *Salvia miltiorrhiza* | 499,236bp | 44.39% | NC_023209 |
| *Rhazya stricta* | 548,608bp | 43.68% | NC_024293 |
| *Capsicum annuum* | 511,530bp | 44.52% | NC_024624 |
| *Hyoscyamus niger* | 501,401bp | 45.18% | NC_026515 |
| *Nicotiana sylvestris* | 430,597bp | 44.96% | NC_029805 |
| *Hesperelaea palmeri* | 658,522bp | 44.47% | NC_031323 |
| *Platycodon grandiflorus* | 1,249,593bp | 43.89% | NC_035958 |
| *Bupleurum falcatum* | 463,792bp | 45.19% | NC_035962 |
| *Nicotiana attenuata* | 394,341bp | 45.05% | NC_036467 |
| *Chrysanthemum boreale* | 211,002bp | 45.36% | NC_039757 |

**Table S4. Species of 60 higher plant mitochondrial genome utilized in the fourth example dataset and their corresponding accession No. in GenBank database.**

| **Species** | **Length** | **GC content** | **Accession NO.** |
| --- | --- | --- | --- |
| *Phoenix dactylifera* | 158,462bp | 37.23% | NC_013991.2 |
| *Elaeis guineensis* | 156,973bp | 37.40% | NC_017602.1 |
| *Elodea canadensis* | 156,700bp | 36.96% | NC_018541.1 |
| *Zingiber spectabile* | 155,890bp | 36.29% | NC_020363.1 |
| *Bismarckia nobilis* | 158,210bp | 37.47% | NC_020366.1 |
| *Cymbidium tortisepalum* | 155,627bp | 37.03% | NC_021431.1 |
| *Cymbidium tracyanum* | 156,286bp | 36.80% | NC_021432.1 |
| *Najas flexilis* | 156,366bp | 38.22% | NC_021936.1 |
| *Cocos nucifera* | 154,731bp | 37.44% | NC_022417.1 |
| *Curcuma roscoeana* | 159,512bp | 36.33% | NC_022928.1 |
| *Magnolia kobus* | 159,443bp | 39.27% | NC_023237.1 |
| *Magnolia liliiflora* | 158,177bp | 39.15% | NC_023238.1 |
| *Fritillaria taipaiensis* | 151,693bp | 36.97% | NC_023247.1 |
| *Dendrobium officinale* | 152,221bp | 37.47% | NC_024019.1 |
| *Dioscorea rotundata* | 155,406bp | 37.20% | NC_024170.1 |
| *Fritillaria cirrhosa* | 151,991bp | 36.95% | NC_024728.1 |
| *Fritillaria hupehensis* | 152,145bp | 36.97% | NC_024736.1 |
| *Bomarea edulis* | 154,925bp | 38.19% | NC_025306.1 |
| *Luzuriaga radicans* | 157,885bp | 38.07% | NC_025333.1 |
| *Corallorhiza macrantha* | 151,031bp | 37.21% | NC_025660.1 |
| *Acorus gramineus* | 152,849bp | 38.70% | NC_026299.1 |
| *Campynema lineare* | 156,261bp | 36.92% | NC_026785.1 |
| *Carludovica palmata* | 158,545bp | 37.74% | NC_026786.1 |
| *Lilium superbum* | 152,069bp | 37.04% | NC_026787.1 |
| *Xerophyllum tenax* | 156,746bp | 37.85% | NC_027158.1 |
| *Heloniopsis tubiflora* | 158,229bp | 37.50% | NC_027159.1 |
| *Trillium cuneatum* | 156,610bp | 37.52% | NC_027185.1 |
| *Trillium decumbens* | 158,552bp | 37.67% | NC_027282.1 |
| *Lilium hansonii* | 152,655bp | 37.00% | NC_027674.1 |
| *Lilium tsingtauense* | 152,710bp | 37.00% | NC_027675.1 |
| *Lilium sp KHK-2014* | 152,715bp | 37.00% | NC_027679.1 |
| *Trillium tschonoskii* | 156,852bp | 37.47% | NC_027739.1 |
| *Colpothrinax cookii* | 157,867bp | 37.30% | NC_028026.1 |
| *Machilus yunnanensis* | 152,622bp | 39.16% | NC_028073.1 |
| *Machilus balansae* | 152,721bp | 39.15% | NC_028074.1 |
| *Phragmipedium longifolium* | 151,157bp | 36.07% | NC_028149.1 |
| *Iris sanguinea* | 152,408bp | 38.05% | NC_029227.1 |
| *Syagrus coronata* | 155,053bp | 37.46% | NC_029241.1 |
| *Goodyera velutina* | 152,692bp | 36.91% | NC_029365.1 |
| *Tofieldia thibetica* | 155,512bp | 37.44% | NC_029813.1 |
| *Potamogeton perfoliatus* | 156,226bp | 36.46% | NC_029814.1 |
| *Lilium distichum* | 152,598bp | 37.06% | NC_029937.1 |
| *Mauritia flexuosa* | 156,367bp | 37.45% | NC_029947.1 |
| *Veitchia arecina* | 157,170bp | 37.47% | NC_029950.1 |
| *Trithrinax brasiliensis* | 158,487bp | 37.33% | NC_029951.1 |
| *Serenoa repens* | 158,952bp | 37.26% | NC_029953.1 |
| *Salacca ramosiana* | 157,047bp | 37.39% | NC_029954.1 |
| *Pritchardia thurstonii* | 157,909bp | 37.28% | NC_029955.1 |
| *Phytelephas aequatorialis* | 159,075bp | 37.23% | NC_029957.1 |
| *Nypa fruticans* | 158,391bp | 37.16% | NC_029958.1 |
| *Lodoicea maldivica* | 159,010bp | 37.32% | NC_029960.1 |
| *Leucothrinax morrisii* | 158,452bp | 37.32% | NC_029961.1 |
| *Eugeissona tristis* | 155,304bp | 37.66% | NC_029963.1 |
| *Chamaerops humilis* | 158,653bp | 37.18% | NC_029967.1 |
| *Brahea brandegeei* | 158,733bp | 37.20% | NC_029968.1 |
| *Borassodendron machadonis* | 158,144bp | 37.44% | NC_029969.1 |
| *Baxteria australis* | 158,858bp | 37.17% | NC_029970.1 |
| *Acoelorraphe wrightii* | 158,503bp | 37.31% | NC_029973.1 |
| *Colchicum autumnale* | 156,462bp | 37.56% | NC_030064.1 |
| *Gloriosa superba* | 157,924bp | 37.59% | NC_030065.1 |

**Table S5. Species of Lamiales chloroplast genomes utilized in the fifth example dataset and their corresponding accession No. in GenBank database.**

| **Species** | **Length** | **GC content** | **Accession NO.** |
| --- | --- | --- | --- |
| *Olea europaeasubsp.europaea* | 155,875bp | 37.81% | FN997650.2 |
| *Sesamum indicum* | 153,324bp | 38.20% | JN637766.2 |
| *Salvia miltiorrhiza* | 151,332bp | 38.02% | HF586694.1 |
| *Scrophularia takesimensis* | 152,425bp | 38.05% | KM590983.1 |
| *Salvia rosmarinus* | 152,462bp | 37.99% | KR232566.1 |
| *Paulownia tomentosa* | 154,540bp | 37.99% | KP718624.1 |
| *Abeliophyllum distichum* | 155,982bp | 37.82% | KT274029.1 |
| *Lavandula angustifolia* | 153,448bp | 38.04% | KT948988.1 |
| *Pedicularisi shidoyana* | 152,571bp | 38.09% | KU170194.1 |
| *Erythranthe lutea* | 153,150bp | 37.72% | KU705476.1 |
| *Castilleja paramensis* | 152,926bp | 38.19% | KT959111.1 |
| *Rehmannia henryi* | 153,890bp | 37.95% | KX636158.1 |
| *Rehmannia solanifolia* | 153,989bp | 37.94% | KX636159.1 |
| *Aloysia citrodora* | 154,699bp | 39.19% | KY085903.1 |
| *Ocimum basilicum* | 152,407bp | 37.84% | KY623639.1 |
| *Forsythia suspensa* | 156,404bp | 37.79% | MF579702.1 |
| *Scrophularia dentata* | 152,600bp | 37.96% | MF861202.1 |
| *Scrophularia henryi* | 152,868bp | 38.00% | MF861203.1 |
| *Chionanthus parkinsonii* | 155,436bp | 37.81% | MG255752.1 |
| *Forsythia xintermedia* | 156,376bp | 37.80% | MG255756.1 |
| *Nestegis apetala* | 154,849bp | 37.83% | MG255758.1 |
| *Olea exasperata* | 155,897bp | 37.81% | MG255766.1 |
| *Pedicularis hallaisanensis* | 143,469bp | 38.65% | NC_037433.1 |
| *Lancea tibetica* | 153,665bp | 37.89% | NC_037693.1 |
| *Salvia officinalis* | 151,089bp | 38.04% | MG772529.1 |
| *Salvia chanryoenica* | 151,689bp | 37.95% | NC_040121.1 |
| *Fraxinus mandshurica* | 155,530bp | 37.82% | MH674342.1 |
| *Salvia przewalskii* | 151,319bp | 37.96% | NC_041091.1 |
| *Salvia bulleyana* | 151,547bp | 37.99% | NC_041092.1 |
| *Schreberatrichoclada* | 155,693bp | 37.77% | MH817942.1 |
| *Buddleja colvilei* | 154,225bp | 38.07% | MH411147.1 |
| *Buddleja sessilifolia* | 154,202bp | 38.10% | MH411149.1 |
| *Forestiera pubescensvar.parviflora* | 155,195bp | 37.90% | MH817905.1. |
| *Schrebera orientalis* | 155,718bp | 37.82% | NC_042266.1 |
| *Schrebera trichoclada* | 155,644bp | 37.80% | NC_042268.1 |
| *Priogymnanthus hasslerianus* | 156,760bp | 37.63% | NC_042279.1 |
| *Chionanthusligustrinus* | 155,912bp | 37.82% | NC_042284.1 |
| *Phaseolusvulgaris* | 155,622bp | 37.82% | MH553374.1 |
| *Brandisiaswinglei* | 155,344bp | 38.07% | MK381315.1 |
| *Wightiaspeciosissima* | 153,621bp | 37.67% | MK381318.1 |
| *Oleaperrieri* | 155,912bp | 37.80% | NC_042375.1 |
| *Schreberacapuronii* | 155,705bp | 37.82% | NC_042388.1 |
| *Fraxinuslanuginosa* | 155,628bp | 37.86% | NC_042424.1 |
| *Picconiaazorica* | 155,350bp | 37.78% | NC_042428.1 |
| *Comoranthusminor* | 155,929bp | 37.79% | NC_042448.1 |
| *Fraxinusquadrangulata* | 155,541bp | 37.84% | NC_042451.1 |
| *Haenianthussalicifolius* | 155,928bp | 37.69% | NC_042453.1 |
| *Nestegiscunninghamii* | 154,907bp | 37.82% | NC_042455.1 |
| *Phillyreaangustifolia* | 155,335bp | 37.81% | NC_042464.1 |
| *Phillyrealatifolia* | 155,300bp | 37.81% | NC_042465.1 |
| *Schreberaalata* | 156,087bp | 37.83% | NC_042467.1 |
